# Analysis of clinical *Candida parapsilosis* isolates reveals copy number variation in key fluconazole resistance genes

**DOI:** 10.1101/2023.12.13.571446

**Authors:** Sean Bergin, Laura A. Doorley, Jeffrey M. Rybak, Kenneth H. Wolfe, Geraldine Butler, Christina A. Cuomo, P. David Rogers

**Author notes:** Joint corresponding authors.

## Abstract

We used whole-genome sequencing to analyse a collection of 35 fluconazole resistant and 7 susceptible *Candida parapsilosis* isolates together with coverage analysis and GWAS techniques to identify new mechanisms of fluconazole resistance. Phylogenetic analysis shows that although the collection is diverse, two probable outbreak groups were identified. We identified copy number variation of two genes, *ERG11* and *CDR1B*, in resistant isolates. Two strains have a CNV at the *ERG11* locus; the entire ORF is amplified in one, and only the promoter region is amplified in the other. We show the annotated telomeric gene *CDR1B* is actually an artefactual *in silico* fusion of two highly similar neighbouring *CDR* genes due to an assembly error in the *C. parapsilosis* CDC317 reference genome. We report highly variable copy numbers of the *CDR1B* region across the collection. Several strains have increased expansion of the two genes into a tandem array of new chimeric genes. Other strains have experienced a deletion between the two genes creating a single gene with a reciprocal chimerism. We find translocations, duplications, and gene conversion across the *CDR* gene family in the *C. parapsilosis* species complex, showing that it is a highly dynamic family.

## Introduction

*Candida parapsilosis* is a human fungal pathogen that is globally one of the most common sources of non-*albicans Candida* infections (1, 2). In the decade 2006-2016, *C. parapsilosis* accounted for ∼16% of all candidemia cases (3). Traditionally, *C. parapsilosis* was predominantly found in immunocompromised patients such as transplant recipients or preterm neonates (4, 5). More recently, however, cases have seen a rise in adult patients in non-surgical wards (6, 7). *C. parapsilosis*, and its sister species *Candida orthopsilosis* and *Candida metapsilosis*, belong to the CUG-Ser1 clade along with other major fungal pathogens *Candida albicans*, *Candida dubliniensis*, and *Candida tropicalis* (8). Unlike its sister species and other members of this clade, *C. parapsilosis* is assumed to be completely asexual due to its high homozygosity, pseudogenisation of *MAT***a**, and the lack of a *MAT*α idiomorph (9, 10). Outbreaks of *C. parapsilosis* have been associated with variants conferring resistance to common antifungal drugs, including fluconazole, a triazole (11). Fluconazole binds to the enzyme lanosterol 14alpha-demethylase, encoded by the gene *ERG11*. This enzyme plays a key role in the ergosterol biosynthesis pathway, which is inhibited by the binding of fluconazole (12–14). Ergosterol is a key component of the fungal cell membrane and in its absence, and with accumulation of alternate sterols, cell growth is arrested (15, 16). Resistance to fluconazole treatment is a growing trend in clinical *Candida spp.* isolates (17). In *C. parapsilosis* resistance is particularly associated with the Y132F substitution in *ERG11* that contributes directly to resistance (18) and has been implicated in many fluconazole-resistant outbreak events across the world (19–24). Equivalent mutations also contribute to fluconazole resistance in *C. albicans*, *C. tropicalis,* and *Candida auris* (25–29).

Most of our understanding of other mechanisms of fluconazole resistance in *Candida* species, including the role of other substitutions in *ERG11*, comes from studies in *C. albicans* (13, 14, 26, 29). Overexpression of *ERG11*, often by gain-of-function mutations in the transcriptional regulator *UPC2*, has been implicated in resistance in *C. albicans* (30–32). In addition, azole resistance is due in part to overexpression of drug efflux pumps (33–35). In *C. albicans*, the contribution of two drug efflux pumps encoded by *CDR1* and *CDR2* that both belong to the ABC transporter (CDR) family to fluconazole resistance has been well studied (33). In the absence of drugs *CDR1* is expressed while *CDR2* is not (36). However, expression of both genes is upregulated in some resistant strains due to activating mutations in *TAC1*, a gene encoding a transcriptional regulator (18, 37, 38). Likewise, overexpression of *MDR1*, which encodes a transporter of the Major Facilitator Superfamily, is overexpressed in some isolates due to activating mutations in *MRR1*, encoding another transcriptional regulator (39). Overexpression of homologs of *CDR1* and *MDR1* have also been found to contribute to resistance in some *C. parapsilosis* clinical isolates that contain similar activating mutations in *TAC1* and *MRR1* (18, 40, 41). Often, multiple resistance mechanisms are found to act in concert in the same isolate leading to high level resistance (42, 43).

In this study we investigate the genetic mechanisms underlying fluconazole resistance in 42 *C. parapsilosis* isolates. Fluconazole resistance has previously been studied in 34 of these isolates using targeted gene sequencing and gene expression analysis (18, 40, 41). Mutations in *ERG11* and over-expression of drug transporters were identified in some isolates. However, some isolates that share the same azole resistance-associated mutation exhibit a range of MIC values, and for other isolates, no obvious resistance mechanisms were identified. Here we use whole genome sequencing, coverage analysis and GWAS methods to identify point mutations and copy number variants (CNVs) associated with novel mechanisms of fluconazole resistance. Using phylogenomic methods, we also identify two probable outbreak clades, from Bloemfontein and Johannesburg, South Africa.

## Results

### Phylogeny and two outbreak clades

Azole resistance mechanisms have previously been studied in 34 fluconazole resistant (MIC ≥ 8 μg/ml) and three fluconazole sensitive (MIC ≤ 2 μg/ml) isolates of *C. parapsilosis* included in this study (18, 40, 41). To improve the power of the analysis (especially for GWAS), we sequenced all 37 genomes, and included one more resistant isolate CDC317 (the reference strain for *C. parapsilosis*) and four susceptible isolates 73/037, 73/114, FM16 and MSK809 (44). The isolates originate from several geographical locations, including several collected from two cities in South Africa between 2001 and 2009 (Johannesburg and Bloemfontein, Table 1). Phylogenomic analysis shows that the isolates represent a broad range of the *C. parapsilosis* phylogeny, as seen when integrated into a tree containing >200 other strains (Fig. S1) (44). Resistant isolates fall into each of the five global clades of *C. parapsilosis* that we have previously identified (44), and susceptible isolates belong to four out of five clades. Despite this breadth, two groups of isolates have very shallow branches, indicating that they have a very close relationship (Fig. 1). The clade marked by a single asterisk contains isolates all originating from the same clinic in Bloemfontein (Table 1). For the clade marked with double asterisks, one isolate comes from Ann Arbor, Michigan whereas the rest originate from a clinic in Johannesburg, collected over a period of eight years. These clades indicate putative outbreaks in the South African clinics.

**Table 1.**
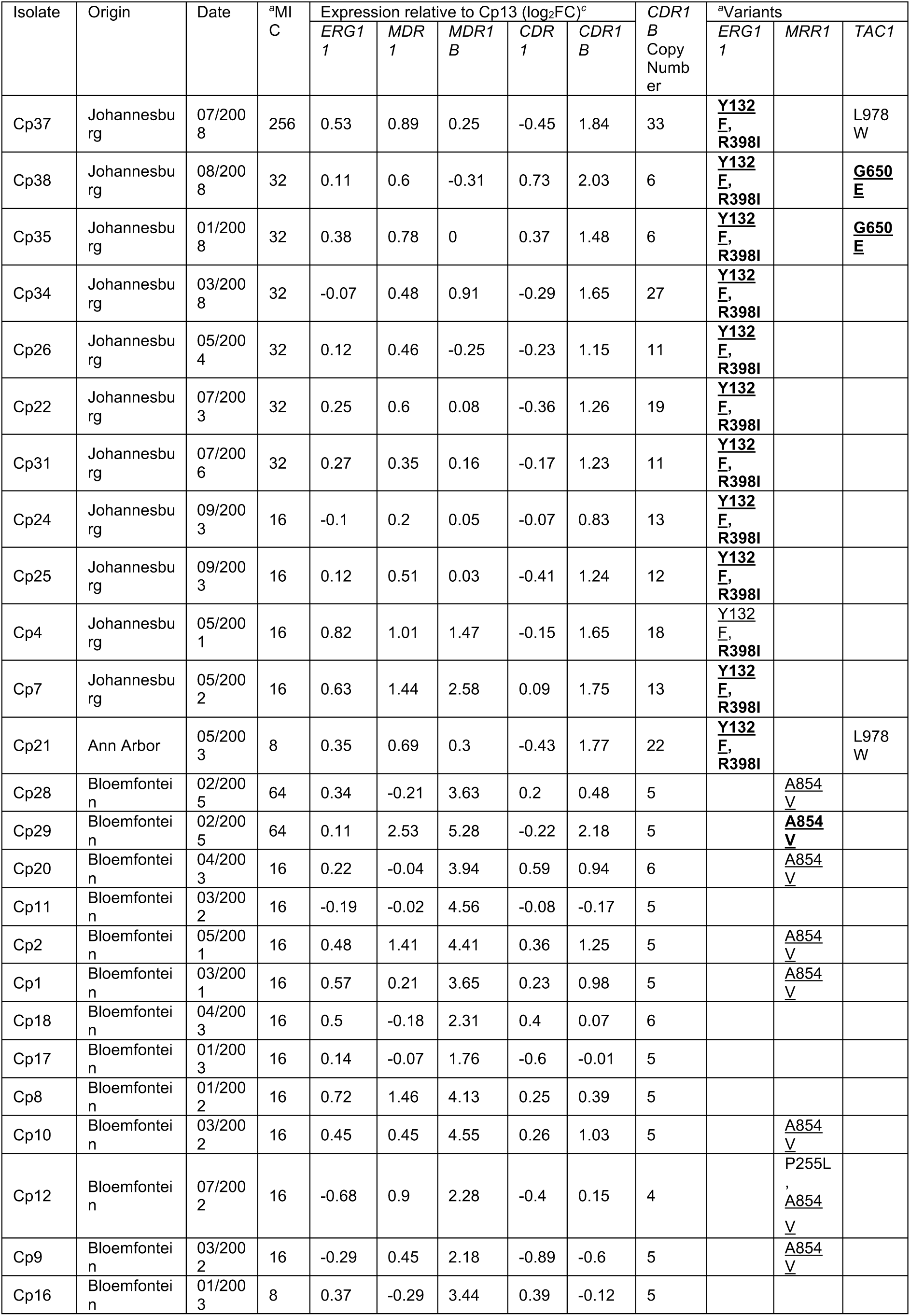
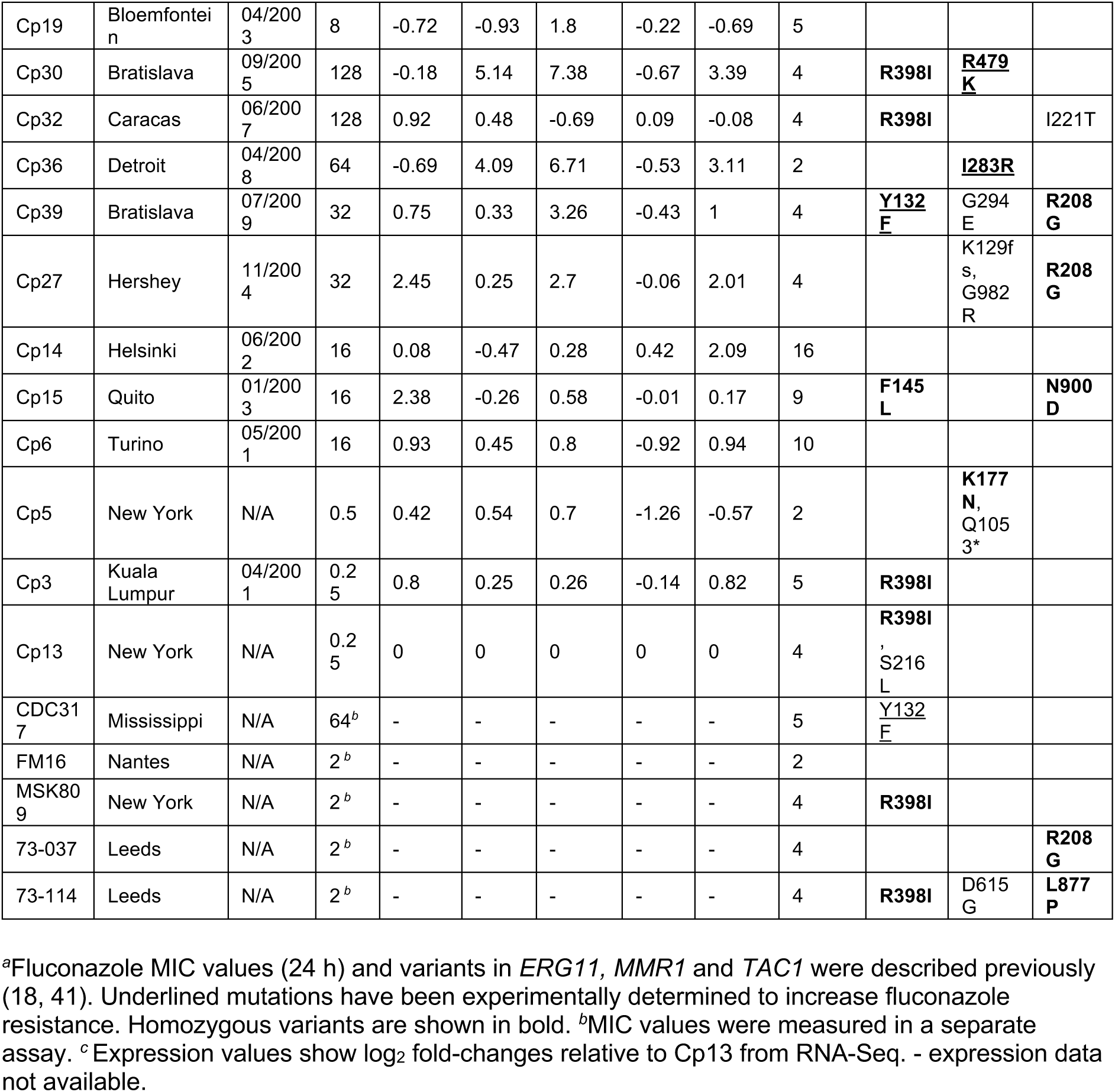
Expression and variant analysis of *C. parapsilosis* isolates.

**Figure 1.**
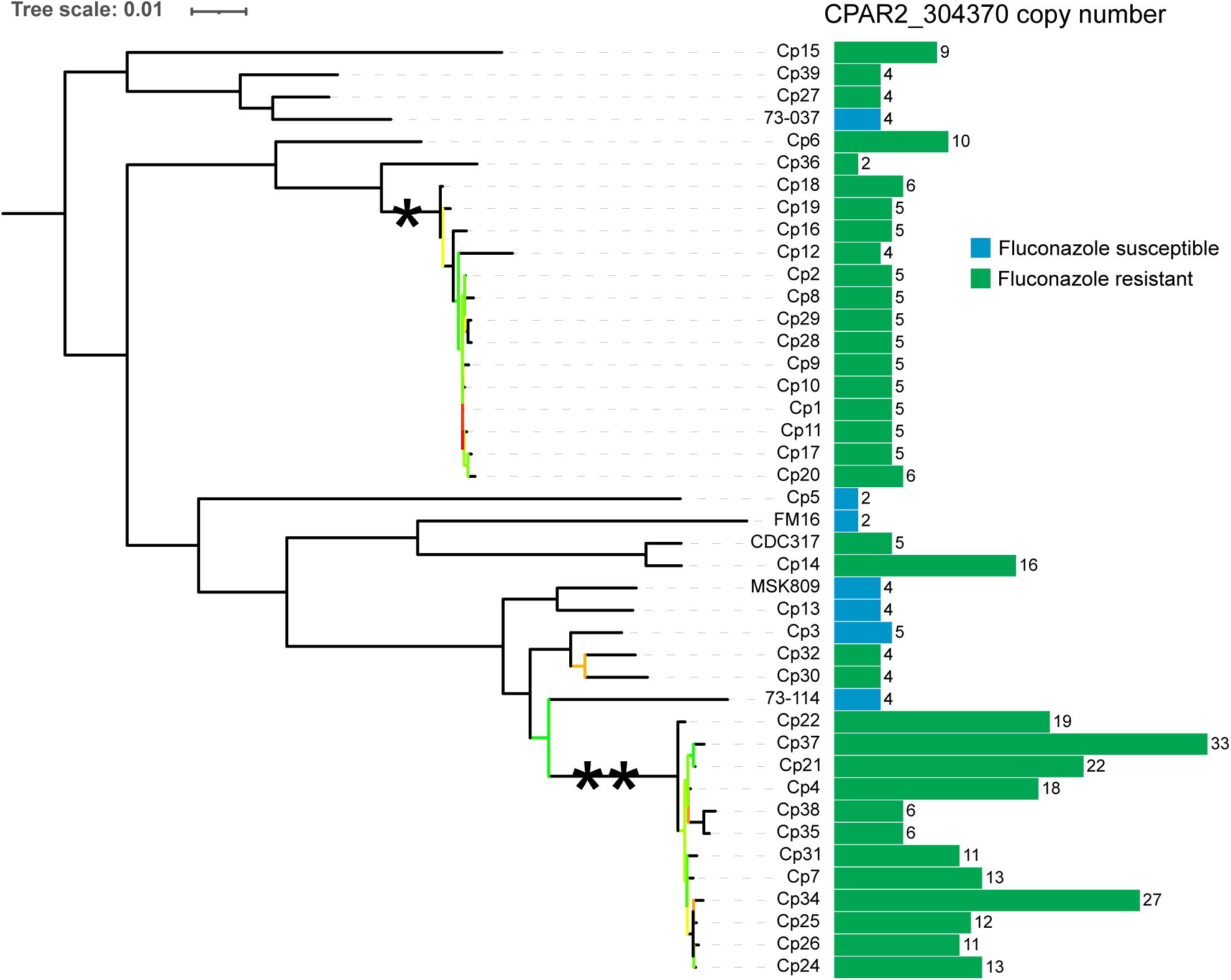
*Left*: Maximum-likelihood tree of 42 *Candida parapsilosis* isolates. RAxML (71) was used to construct the tree from an alignment of 15,582 SNPs genome-wide, using the GTRGAMMA model of nucleotide substitution. Branches with bootstrap values < 100 after 1000 iterations of bootstrap sampling are coloured according to value, ranging from red (0) to green (99). Clades are marked with asterisks to denote potential outbreaks in Bloemfontein (*) and Johannesburg (**). Right: Bar chart showing estimated copy number of the *CDR1B* locus (*CPAR2_304370* in the reference genome assembly). Fluconazole-susceptible isolates are denoted with a blue bar, resistant isolates with a green bar. Copy number was estimated by taking mean coverage across the *CPAR2_304370* ORF and dividing by half the modal cover-age for the isolate.

For 34 of the fluconazole-resistant isolates, multiple potential resistance mechanisms were previously identified using gene expression analysis (RT-qPCR) and targeted gene sequencing (18, 40, 41) (Table 1). To improve the resolution we measured the expression of target genes using RNA-seq, by comparing to the expression of the genes in the azole susceptible isolate Cp13 (Table 1). The RNA-seq data supports the previous analysis (40, 41). For example, overexpression of drug transporter *CPAR2_603010* (*MDR1B*) in strains from Bloemfontein (log_2_FC ranging from 1.8 to 5.3, Table 1), mediated at least in part by the A854V activating mutation in the regulator gene *CPAR2_807270* (*MRR1*), contributes directly to resistance to fluconazole (41). Some Bloemfontein strains have increased expression of *MDR1B* but do not have a corresponding *MRR1* mutation (e.g. Cp11, Table 1) (41). Strikingly, the isolate homozygous for A854V, Cp29, has much higher expression of both *MDR1B* (log_2_FC = 5.28) and *CDR1B* (log_2_FC = 2.18) compared to the other Bloemfontein strains. In addition, mutations in the ergosterol-biosynthesis gene *CPAR2_303740* (*ERG11*) and the *CDR*-family regulator *CPAR2_303510* (*TAC1*) were shown to contribute to fluconazole resistance in other isolates (40).

Fourteen strains (including the reference strain *C. parapsilosis* CDC317) harbour the *ERG11* Y132F substitution which is a well-documented resistance mutation (11, 17, 18, 25, 45). The Y132F substitution is heterozygous in CDC317 and in Cp4, and homozygous in the other 12. The isolates from Johannesburg all have the Y132F mutation (including the heterozygous Cp4), but the isolates from Bloemfontein do not (Table 1).

Eight of the 35 resistant strains do not have any mutations in *ERG11*, *TAC1*, or *MRR1* that have been experimentally determined to affect fluconazole resistance (Table 1). The R398I mutation in *ERG11* has been frequently observed occurring in tandem with Y132F (21, 23), but has also been found without Y132F in susceptible isolates (40). The substitutions A854V, R479K, and I283R in *MRR1* have all been identified as activating mutations leading to the upregulation of genes including *CDR1B,* and *MDR1B* (46). The *TAC1* G650E mutation has shown to increase fluconazole resistance and overexpress *CDR1* and *CDR1B* when introduced into a susceptible isolate (18). In addition, among this collection of isolates, strains that share the same mutation can differ 32-fold in their MIC values (e.g. isolates in the Johannesburg clade have MICs varying from 8 to 256 μg/ml, Table 1). This suggests that novel resistance mechanisms remain to be identified, and that different mechanisms may have additive effects that have not been captured by these studies.

### Copy number variation of *ERG11*

We found that aneuploidy is relatively common in the 42 isolates; 13 have either segmental or whole chromosome aneuploidies (Fig. S2), and several isolates have aneuploidies of multiple chromosomes. However, unlike in *C. albicans* where aneuploidy of the chromosome containing *ERG11* and *TAC1* has been associated with fluconazole resistance (47–49), none of the resistant strains in the collection have extra copies of chromosome 3. Several resistant strains have aneuploidy of chromosome 5, which contains the *ERG4* gene. The potential role of *ERG4* in fluconazole resistance in *Candida spp.* has not been well characterised, but the gene has been found overexpressed alongside *ERG11* in azole-resistant *C. albicans* (50).

We found that two resistant strains, Cp15 and Cp27, have small CNVs at the *ERG11* locus on chromosome 3 (Fig. 2). In Cp27, the entire coding sequence of both *ERG11* and the upstream gene (*HMS1*), and part of the downstream gene (*THR1*), has been amplified by the CNV (1,309,908-1,315,502 bp). Here, the locus has been amplified to nine copies. Short read mate-pair mapping supports the interpretation that this CNV is a tandem array of duplicated sequence. In Cp15, a 341 bp section of the *ERG11* promoter region is amplified to eight copies (1,312,556-1,312,896 bp). Cp15 and Cp27 are the only two isolates in this collection with increased expression of *ERG11* (Table 1). As far as we are aware, this is the first time that an amplification of an *ERG11* promoter has been observed in any *Candida* species.

**Figure 2.**
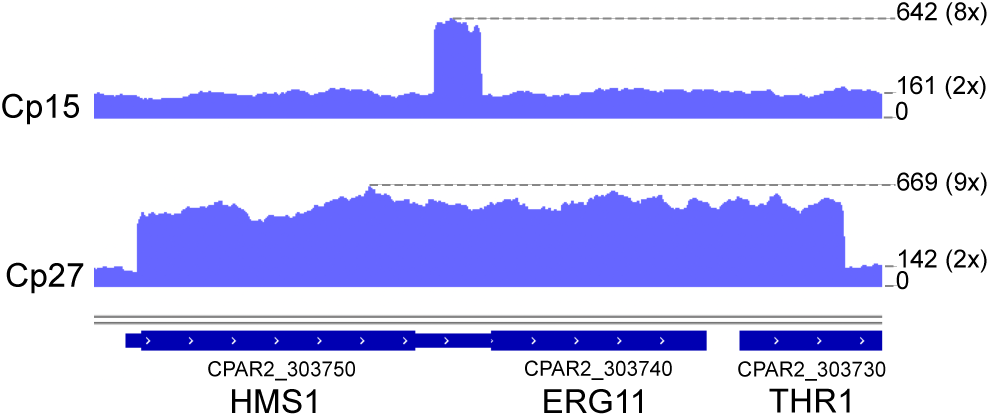
Amplification of *ERG11* in fluconazole resistant isolates *C. parapsilosis* Cp15 and Cp27. Screenshot from IGV showing coverage (read depth) tracks (75). Coverage value is marked on the right for the peak of the amplification, and for the region average. Estimated copy numbers are in parentheses. Thick bars at bottom show CDS of genes, and thinner bars represent transcribed regions.

### Genome Wide Association Study

We performed a Genome Wide Association Study (GWAS) using all 42 isolates to identify potential variants associated with fluconazole resistance that had not been found in earlier studies. The GWAS was carried out using GEMMA (Genome-wide Efficient Mixed Model Association) (51), which calculates and incorporates relatedness data between isolates in order to minimise the confounding effect of population structure on association scores. Because the MIC assays for fluconazole were measured in two different laboratories (Table 1), the phenotypic data was converted to a binary score of either resistant or susceptible to reduce possible bias, with MIC ≤ 2 μg/ml classed as susceptible and MIC ≥ 8 μg/ml classed as resistant, in line with CLSI (52) and EUCAST (53) guidelines. Genotypes were also converted into binary presence/absence of non-reference alleles, with both heterozygous and homozygous variants treated as present. In addition, only variants that were predicted to affect protein function by SIFT (54) were included in the analysis to narrow the search. In total, 7462 variants were used as input to GEMMA. The GWAS analysis did not identify any significant associations below the Bonferroni-corrected p-value threshold of 6.7 x 10^-6^ (Table S1). However, several of the variants with the lowest p-value scores were in *CPAR2_405290* (*CDR1*) and *CPAR2_304370* (*CDR1B*), members of the ABC family of putative drug transporters. Investigating the alignments leading to these calls showed that the *CPAR2_405290* variants are likely a result of mismapping from a similar gene in the genome, so we did not investigate *CPAR2_405290* further. Overexpression of *CPAR2_304370* (*CDR1B*) has previously been observed in fluconazole-resistant isolates and shown to directly contribute to this phenotype (40, 41, 46). In a study investigating acquired azole resistance in consecutive isolates taken from a patient undergoing fluconazole treatment, one isolate with reduced susceptibility to fluconazole had undergone amplification of the *CDR1B* locus, from 4 to ∼15 copies (46). We therefore investigated the *CDR1B* locus in more depth.

### The *CDR1B* locus contains two genes, *CDR1B.1* and *CDR1B.2*, and is amplified in resistant isolates

While further characterising the role of *CPAR2_304370* (the annotated *CDR1B* gene in the reference genome assembly) in azole resistance, we noticed an increased sequence coverage compared to genomic average in most of the isolates. We used the average coverage across the ORF to estimate the copy number of this gene in each of the isolates. Whereas a majority (25/42) of the isolates have a copy number in the range 4-6x, there are several outliers ranging up to 33x, and only three isolates have the expected value of 2x (Fig. 1). Several isolates that have increased expression of *CDR1B* but no corresponding *MRR1* gain-of-function mutation have increased copy number of *CDR1B*, suggesting that amplification of this locus and activating mutations in upstream regulators can both drive overexpression of this gene. Strikingly two of the three isolates with two copies of the locus are susceptible to fluconazole, and no susceptible isolate has more than five copies.

Of special note is the clade containing *C. parapsilosis* FM16, CDC317, and Cp14 (Fig. 1). FM16 is susceptible to fluconazole and has only two copies of *CPAR2_304370*. CDC317 and Cp14 are both fluconazole resistant. CDC317 has only 5 copies of *CPAR2_304370*. However, CDC317 is heterozygous for a Y132F mutation in *ERG11* that is not present in Cp14 (Table 1). In contrast, Cp14 has 16 copies of *CPAR2_304370*. We propose that the related isolates Cp14 and CDC317 acquired resistance by differing mechanisms, the former by acquiring a mutation in *ERG11*, and the latter through increased copy number of *CPAR2_304370*.

The MIC values of the isolates from the Johannesburg outbreak range from 16-256 μg/ml (18) (Table 1). Two related isolates (Cp38 and Cp35) with MICs of 32 μg/ml have acquired a G650E substitution in *TAC1*, resulting in increased expression of *CDR1* (Table 1) (18). The combination of Y132F in *ERG11* and G650E in *TAC1* likely increases resistance compared to each variant alone (18). The copy number of *CDR1B* is highly variable in the Johannesburg isolates, ranging from 6-33x (Fig. 1). Whereas there is no direct correlation between the copy number of *CDR1B* and MIC in these isolates, it is notable that Cp37 has the highest MIC (256 μg/ml) and the highest number of *CDR1B* copies (∼33).

Long-read (Oxford Nanopore) sequencing of CDC317 revealed that the *CPAR2_304370* gene annotated in the Sanger sequencing reference genome assembly of this strain was in fact erroneously assembled by fusing together two highly similar tandem genes, which we now call *CDR1B.1* and *CDR1B.2* (Fig. 3A). As a result, the intergenic space between these two genes, and parts of the genes themselves, are not present in the original reference assembly. It is likely that the presence of *CDR1B.1* and *CDR1B.2* is the ancestral (and most common) state of the locus in *C. parapsilosis* (Fig. 3A), and that the four copies of *CPAR2_304370* indicated in many of the isolates by coverage analysis relative to the reference genome assembly in fact represent diploids with two copies of both *CDR1B.1* and *CDR1B.2* (Fig. 1). We used the long sequencing reads, alongside short reads, to generate a highly accurate, complete chromosome assembly of CDC317. This assembly confirmed that CDC317 has two copies each of *CDR1B.1* and *CDR1B.2*, so the previously estimated copy number of 5x *CPAR2_304370* (Fig. 1) was likely inflated by short reads mismapping from related genes. Using the new accurate assembly of CDC317, we found that *CDR1B.1* and *CDR1B.2* are 98.69% identical at the nucleotide level and the intergenic regions upstream of both genes are 46.10% identical. The two genes differ by only 23 amino acids (out of 1498) when translated.

**Figure 3.**
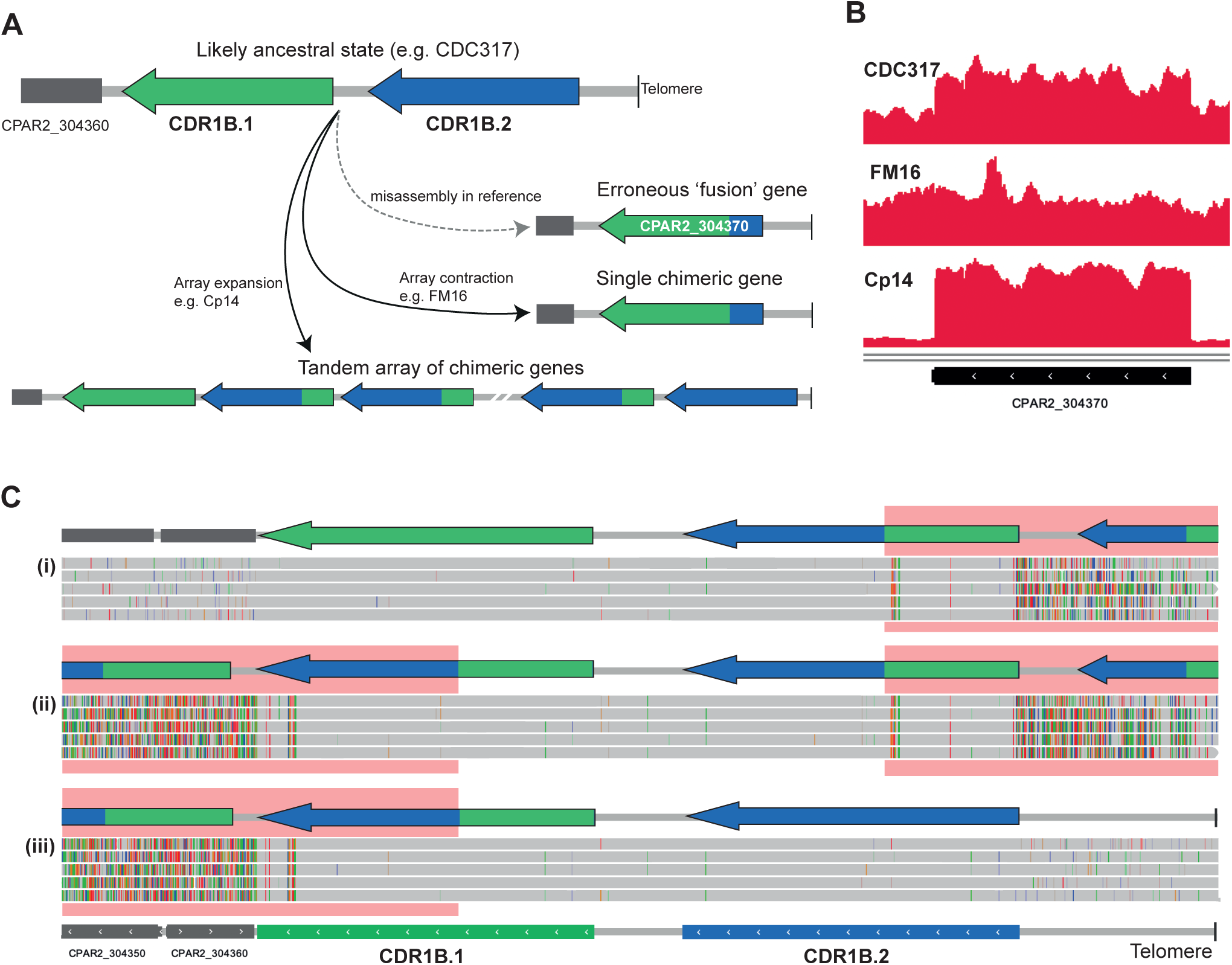
A. Diagram showing the likely ancestral state of the *CDR1B* (*CPAR2_304370*) locus, with two highly similar genes *CDR1B.1* and *CDR1B.2* in tandem as occurs in the corrected genome sequence of C. parapsilosis CDC317 (based on MinION sequencing). The dashed grey arrow indicates how mis-assembly of this locus in the original reference genome sequence for CDC317 (76) led to an erroneous fusion between *CDR1B.1* and *CDR1B.2*, resulting in annotation of the incorrect gene structure *CPAR2_304370*. The solid grey arrows show CNVs formed by array expansion and or contraction in other isolates, such as Cp14 and FM16. B. Coverage tracks in IGV of short-read data from three isolates aligned to the original *C. parapsilosis* CDC317 assembly. Both CDC317 and Cp14 have increased coverage at the *CPAR2_304370* locus, whereas FM16 does not. C. MinION reads of isolate Cp14 aligned to a long-read CDC317 assembly show evidence of a tandem array of chimeric genes bounded by parental genes. The bottom track shows the annotated long-read assembly of CDC317, containing both *CDR1B.1* and *CDR1B.2.* Each grey bar is a single read aligned to the *CDR1B* locus where grey denotes read sequence matching the reference, and coloured dashes are mismatched positions, from modified screenshots of IGV (70). Reads aligned to this locus belong to one of three groups: (i) reads that align fully to *CDR1B.1* and left flanking DNA while partially aligning to CDR1B.2, (ii) reads that align partially to both *CDR1B.1* and *CDR1B.2*, and (iii) reads that align fully to CDR1B.2 and right flanking DNA while partially aligning to *CDR1B.1*. A schematic showing the genic content for each read group is shown above for clarity. Red boxes highlight regions where the read sequence does not match the reference sequence. Reads from groups (i) and (iii) contain one parent gene and one or more copies of the tandem chimera. Reads from group (ii) contain multiple copies of the tandem chimera but neither of the parent genes.

Significantly, short reads from isolate FM16 (estimated to have two copies of *CPAR2_304370*) map to the original *C. parapsilosis* CDC317 reference genome without an increase in coverage or misaligned read pairs (Fig. 3B). This is evidence of an array contraction of the two genes in FM16 that results in a single chimeric *CDR1B.2/CDR1B.1* gene, biologically mirroring the misassembly in the original reference. Short read alignments of Cp36 and Cp5 suggest that similar array contractions occurred in these isolates.

Long-read sequencing of Cp14 revealed that the extra copies predicted by coverage analysis (16x *CPAR2_304370*) are the result of a tandem array of identical chimeric *CDR1B.1/CDR1B.2* genes, bounded upstream by non-chimeric *CDR1B.2* and downstream by non-chimeric *CDR1B.1* (Fig. 3A). These chimeric genes inherited their 5’ region from *CDR1B.1* and their 3’ region from *CDR1B.2*. In this manner they are opposite to the chimeric gene in FM16. None of the Cp14 long sequencing reads reached across the entire tandem array, so the exact copy number of the chimeric genes could not be determined. However, by aligning the reads to the long-read assembly of CDC317, we found reads that contain the beginning, middle, and end of the tandem array (Fig. 3C).

### Families of *CDR* orthologs and paralogs in the *C. parapsilosis* clade

*CDR1B.1* and *CDR1B.2* are two of nine *CDR* genes in *C. parapsilosis* (Fig. 4). Strikingly, most (5/9) of these genes are located in telomeric regions. Many of the *CDR* genes in *C. parapsilosis*, including all five telomeric ones, have direct orthologs in *C. metapsilosis* and *C. orthopsilosis* but they are more distantly related to *C. albicans CDR* genes. The *CDR* orthologs in *C. metapsilosis* and *C. orthopsilosis* share synteny of neighbouring genes when compared to the gene order of *C. parapsilosis*. The telomeric *CDR* genes are likely to have originated after *C. parapsilosis* diverged from the *C. albicans* lineage, but before the separation of *C. parapsilosis* from *C. orthopsilosis* and *C. metapsilosis*. In addition, a recent gene duplication in *C. parapsilosis* produced the gene pair *CPAR2_300010/CPAR2_603800* which are duplicated only in this species, while the *CDR1B.1/CDR1B.2* gene pair has a more complicated history.

**Figure 4.**
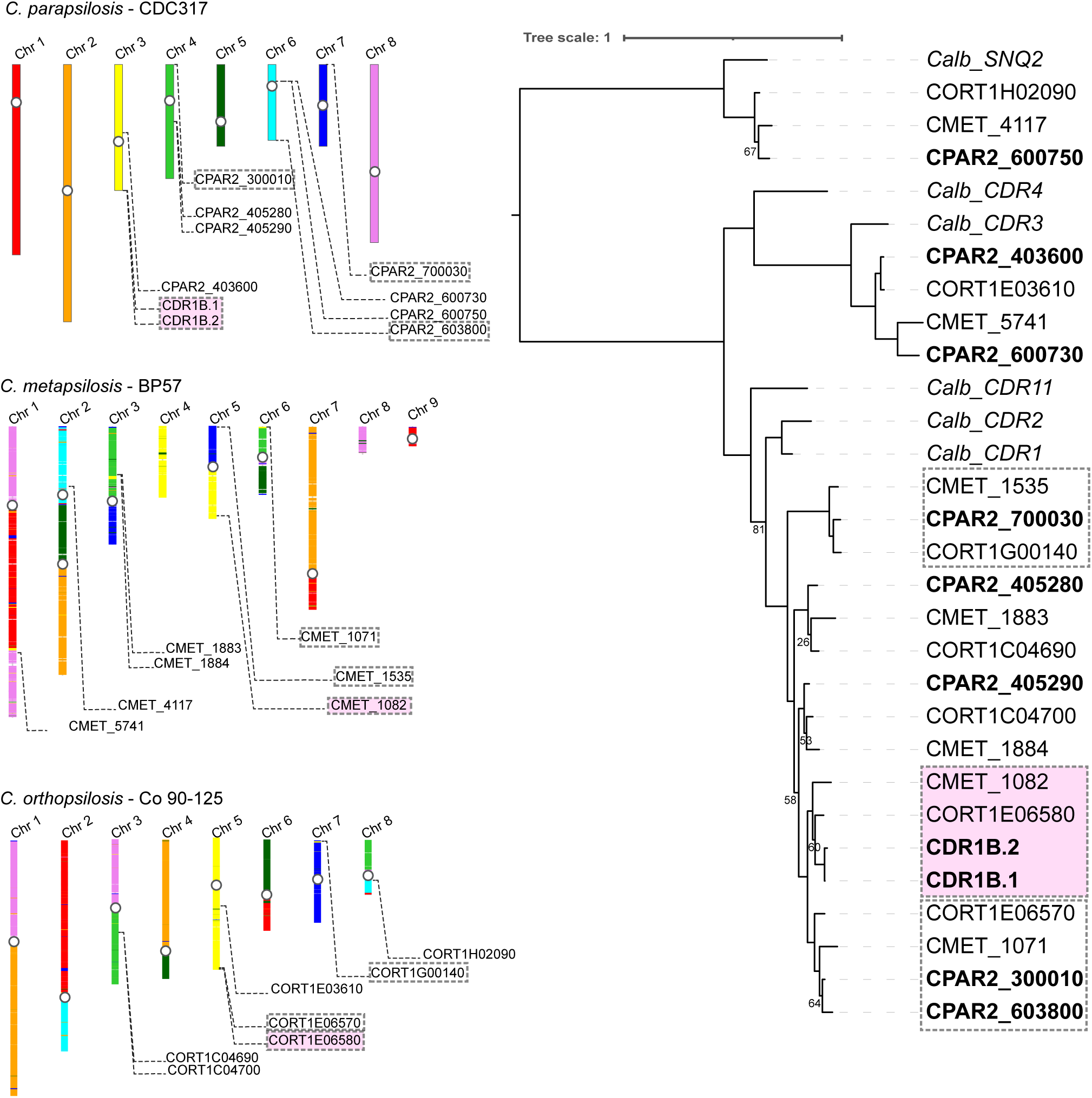
Left: Schematic of synteny shared between members of the *C. parapsilosis* species complex. The *C. parapsilosis* genome was aligned to the genomes of *C. metapsilosis* (GCA_017655625.1) (55) and *C. orthopsilosis* (PRJNA767198) using BLASTN. These genomes were then coloured according to the *C. parapsilosis* chromosome aligned to each region. Previously identified centromeres are marked with a white circle (77). The locations of CDR genes are shown for each species. Right: Phylogenetic tree of CDR protein sequences from *C. parapsilosis*, *C. orthopsilosis*, *C. metapsilosis*, and *C. albican*s. Pink backgrounds denote genes most closely related to *CDR1B.1* and *CDR1B.2*. Three clades of genes located near telomeres are marked with boxes. Protein sequences were aligned using the Clustal Omega method in Seaview (78). The tree was constructed with the LG model within Seaview, using 100 bootstrap replicates. Bootstrap values >85 have been omitted for clarity.

*C. metapsilosis* has only a single gene (*CMET_1082*) at the *CDR1B* locus, which is equally related to the two *C. parapsilosis CDR1B* genes (Fig. 4; genes highlighted in pink). Long-read sequences of the reference *C. metapsilosis* strain BP57, recently assembled by Mixão et al. (52), indicate that this is not due to misassembly. There are two *CDR* genes present at the *CDR1B* locus in *C. orthopsilosis*, but interestingly only one of them (*CORT1E06580*) falls phylogenetically into the same clade as *C. parapsilosis CDR1B.1* and *CDR1B.2* and *C. metapsilosis CMET_1082*, while its neighbour (*CORT1E06570*) falls into an adjacent clade with *CMET_1071* and two *C. parapsilosis* genes *CPAR2_300010* and *CPAR2_603800* (Fig. 4). Notably, the *CDR1B* locus, *CPAR2_300010* and *CPAR2_603800* are all telomeric on different chromosomes of *C. parapsilosis*, so we suggest there may have been some *CDR* gene exchange and/or homogenization among telomeric regions within these species.

We did not observe CNVs affecting any of the other *CDR* genes in *C. parapsilosis*, nor in 36 *C. orthopsilosis* or 30 *C. metapsilosis* isolates that we analyzed. The *CDR1B* locus of *C. parapsilosis* is unique among the *CDR* genes of these three species in having two highly similar genes in tandem, which provides a template for amplification of the locus to occur readily.

## Discussion

Using a genome-wide approach, we identified two CNVs (*ERG11* and *CDR1B.1*/*CDR1B.2*) that are associated with fluconazole resistance in *C. parapsilosis*. CNVs are a method of gene duplication by which an organism can transiently adapt to its environment (53, 54). Environmental changes, such as introduction of an antifungal drug, can select for specific genes to be duplicated, and thereby overexpressed (49, 55). After the drug is removed, the CNV can be lost by selective pressure to maintain a compact genome size (55).

Overexpression of *ERG11* by increasing copy number has been observed in *C. albicans* (47, 48). However, in *C. albicans* the gene was amplified along with *TAC1* by means of a partial aneuploidy of chromosome 5 leading to a formation of an isochromosome, i(5L). The i(5L) isochromosome, which typically results in a single extra copy of the chromosomal region, has been identified in multiple clinical isolates where it has a modest but measurable impact on fluconazole resistance in different genetic backgrounds (47, 49, 56). Genomic expansion of *ERG11* in combination with hotspot mutations is also associated with azole resistance of *C. tropicalis* (57). In *C. auris*, a large survey of 304 isolates identified a CNV including *ERG11* in 18 isolates (most from a single clade) which was associated with fluconazole resistance (58). Recently, a laboratory-directed evolution experiment also showed that reduced azole susceptibility is associated with large segmental duplications containing *ERG11* in *C. auris*. One evolved strain had a 191 kb long CNV with 75 protein encoding genes including *ERG11* amplified, while another had a 161 kb long CNV containing 67 protein encoding genes including *ERG11* (59).

A very recent analysis identified amplifications of *ERG11* in 21 azole resistant isolates of *C. parapsilosis* (60). The amplifications ranged from partial aneuploidy of chromosome 3, similar to *C. albicans* (47, 48), to smaller amplifications of 2.3 to 12.1 kb (60). These are similar to the amplifications that we observe in strain Cp27, where *ERG11* and its neighboring gene are amplified (5.6 kb). The CNV in strain Cp15 is distinctly different; only the *ERG11* promoter region is amplified. Importantly, both Cp27 and Cp15 isolates also have increased *ERG11* expression (logFC ∼2.4, Table 1), strongly suggesting these duplications directly impact expression of this gene and fluconazole resistance. We have previously observed a similar localised gene amplification in *C. parapsilosis* in the gene *RTA3* (44). Several different CNVs, spanning either the whole gene or just the promoter region, led to overexpression of *RTA3* associated with increased resistance to the antimicrobial drug miltefosine. Promoter amplifications may therefore be a previously underexplored mechanism of drug resistance in *Candida* species.

The GWAS analysis performed in this study failed to find significant associations. Although disappointing, it is within expectation because the sample size (n=42) is very low for a study of this kind, and there was a large imbalance between the number of resistant (n=35) and susceptible (n=7) isolates. PowerBacGWAS, a tool used to find required sample sizes for GWAS in bacteria, estimates in a best case scenario where minor allele frequency is high and effect size is large, analysis of 500 isolates would be required to identify a single significant SNP in *Mycobacterium tuberculosis* (61). This issue is further compounded in our study by the presence of groups of highly related strains from outbreak infections, which reduces the effective sample size. Despite the lack of statistical power, the GWAS results guided us towards characterising the *CDR1B* amplification as a possible mechanism of fluconazole resistance. This discovery lends support to the use of GWAS in *C. parapsilosis,* particularly in studying easily definable phenotypes such as antifungal resistance.

Our analysis of the *CDR* family in the *C. parapsilosis* species complex has led to some interesting insights. We found that most of the *CDR* genes in the *C. parapsilosis* species complex have arisen by gene duplication events after the lineage diverged from *C. albicans*. The *CDR* gene content varies between *C. parapsilosis* and its two sister species. Many *CDR* copies are located at telomeres and there is evidence of exchange of duplicated genes between telomeres. The *CDR1B* locus is particularly variable between the three sister species; there is a single gene in *C. metapsilosis*, two distinct genes in *C. orthopsilosis*, and two highly similar genes in *C. parapsilosis*. It is possible that *C. orthopsilosis* represents the ancestral state, with one of the genes lost in the *C. metapsilosis* lineage and one gene overwritten in *C. parapsilosis* by gene conversion from its neighbour.

Previous analysis identified several potential azole resistance mechanisms in some of the strains described here (18, 40, 41). This includes the Y132F mutation in *ERG11* (Table 1) (18). We find that this variant is present in all the closely related isolates from Johannesburg, supporting our inference that these represent an infection outbreak. However, the range of MICs observed in these isolates cannot fully be explained by the presence of Y132F (Table 1). Two isolates (Cp35 and Cp38) collected in 2008, late in the inferred outbreak, have also acquired variants in Tac1, with an associated rise in expression of *CDR1* (Table 1, logFC 0.4-0.7) (18). Our analysis suggests that in other isolates (e.g. Cp37, also collected in 2008) increased resistance is associated with, and may be caused by, amplification of the *CDR1B* locus (up to ∼33 copies).

The presence of *ERG11* Y132F is increasingly associated with infection outbreaks (19–23, 25), suggesting that it may confer a transmission advantage beyond the effect on azole resistance. However, outbreaks can also be caused by strains without Y132F such as the Bloemfontein isolates in this study. For example, in a study investigating 60 *C. parapsilosis* strains involved in a large outbreak in a Brazilian ICU, only ∼36% of isolates resistant to fluconazole had an *ERG11* mutation (62). Another outbreak among patients undergoing allo-hematopoietic cell transplant treatment was associated with isolates without the Y132F mutation (63).

All of strains from the Bloemfontein outbreak have increased expression of *MDR1B* (log_2_FC 1.8-5.28, Table 1). Although the isolates are closely related, there is some variability in their *MRR1* alleles. Some isolates contain an A854V activating mutation in *MRR1*, which is known to result in overexpression of *MDR1B* (41). Seven are heterozygous for the A854V mutation in *MRR1*, one is homozygous for the mutation, and six do not have the mutation. The earliest cultured strains (Cp1 and Cp2 isolated in 2001), are both heterozygous for the mutation whereas some strains isolated later (e.g. Cp17 in 2003) are lacking the mutation entirely (Table 1). This indicates that there may be sub-populations of related strains existing concurrently in the clinic which are variable at *MRR1*. Multiple strains without Mrr1 A854V have increased expression of *MDR1B* (Table 1), showing that there may be additional unidentified factors contributing to *MDR1B* expression in this clade.

Eight of the resistant isolates in Table 1 contain none of the common variants associated with azole resistance. Our work identifies relevant CNVs in 4 of these. *ERG11* is amplified in two strains, Cp27 and Cp15, that are not associated with outbreaks. Strains Cp14 and Cp6, which were also not associated with outbreaks, both have >10 copies of *CDR1B* and no other identified resistance mechanism. The variation in copy number of *CDR1B* across the isolates suggests that *CDR1B* amplification may be a common mechanism of azole resistance in *C. parapsilosis*. It is important to note that, of the multiple *CDR* genes in multiple *Candida* species, we observed CNVs at only the *C. parapsilosis CDR1B* locus. We propose that the existence of two nearly identical genes in tandem makes CNVs at this locus more likely to occur. In this way, *C. parapsilosis* may be primed to generate extra copies of *CDR1B*, and therefore be predisposed to develop fluconazole resistance. Amplification likely occurs during infection as described by Branco et al (46), where a strain of *C. parapsilosis* in a patient treated with fluconazole acquired a CNV amplifying *CDR1B* that was associated with reduced fluconazole susceptibility.

## Methods

### RNA Sequencing

*C. parapsilosis* isolates were maintained at -80°C in 40% glycerol stocks. Isolates were grown in YPD liquid media overnight and plated onto Sabouraud-Dextrose (BD companies) agarose plates in biological triplicate for 24h growth at 30°C. Sterile loops were used to inoculate 20mL RPMI with MOPS and 2% glucose to OD600=0.1. Cultures were incubated at 35°C with110rpm shaking for 8h, after which the cells were centrifuged at 4000rpm for 5 min. Supernatants were removed and pellets were stored at -80°C for a minimum of 24hrs. RNA isolation was performed via RiboPureTM Yeast (Invitrogen) kits per manufacturer’s instructions. RNA Sequencing performed using Illumina NextSeq for stranded mRNA (Hartwell Center, St Jude Children’s Research Hospital). Libraries were prepared with paired-end adapters using Illumina chemistries per manufacturer’s instructions, with read lengths of approximately 150bp with at least 50 million raw reads per sample (Bioproject 14022043). RNA-sequencing was analyzed using CLC Genomics Workbench version 20.0 (QIAGEN), and reads were trimmed using default settings for failed reads and adaptor sequences and then subsequently mapped to the *C. parapsilosis* genome (GenBank accession: GCA_000182765.2) with paired reads counted as one and expression values set to RPKM. Principal-component analysis was utilized for initial assessment of biological replicate clusters. Whole transcriptome differential gene expression analysis was performed with the prescribed algorithm of CLC Genomics Workbench version 20.0. Mismatch, insertion, and deletion costs were set to default parameters and a Wald test was used to compare all isolates against the fluconazole susceptible isolate Cp13. Fold changes for CPAR2_304370, CPAR2_405290, CPAR2_301760 were identified for all isolates and are reported in Table 1.

### Whole Genome Sequencing

Genomic DNA was isolated from overnight yeast peptone dextrose (YPD) liquid media cultures utilizing a Triton SDS and phenol-chloroform method previously described by Amberg et al (64). DNA concentrations were quantified using both the Qubit Fluorometer and Nanodrop spectrophotometer using the manufacturers’ protocols. Whole genome libraries were prepared and sequenced on the NovaSeq600 platform (150 bp, paired-end reads) by the University of Maryland School of Medicine Institute for Genomic Sciences.

### Fluconazole Susceptibility Testing

Inoculums of YPD liquid media were prepared from original stocks, stored at −80°C in 40% glycerol. Inoculates were grown at 30°C with 220 rpm shaking overnight and subsequently plated onto Sabouraud dextrose agar for overnight incubation at 35°C. Minimum inhibitory concentrations (MIC) were determined in RPMI1640 (Roswell Park Memorial Institute) supplemented with MOPS (3-(N-morpholino) propanesulfonic acid) buffer and 2% glucose, pH 7.0 liquid media following CLSI M27-A4 methods for broth microdilution. Fluconazole (Sigma Aldrich) drug stocks were prepared in dimethyl sulfoxide (DMSO) at 100× the maximum plate concentration (256 mg/mL for resistant isolates and 16 mg/mL for susceptible isolates). MICs were determined visually as the concentration achieving 50% growth inhibition at 24 hours, and the modal value of biological triplicate measurements was considered the MIC reporting.

### MinION sequencing

Samples were grown overnight in 50 mL of YPD broth at 30°C. Genomic DNA was extracted from 1.5 ml of liquid cultures saturated to 10 A600 units per millilitre using the Yeast Masterpure DNA purification kit (MPY80010) following manufacturer’s instructions.

Genomic DNA (1 µg) from each sample sequenced with MinION technology using the native barcoding kits (SQK-NBD-24 and SQK-NBD114-24) from Oxford Nanopore Technologies (ONT), following manufacturer’s instructions. Library kit SQK-NBD112-24 was used for sequencing CDC317 on an R9.4.1 chemistry flowcell (FLO-MIN106D). Library kit SQK-NBD114-24 was used for sequencing Cp14 on an R10.4.1 chemistry flowcell (FLO-MIN114). Sequencing of both strains was performed on a MinION MK1C device with MinKNOW (ONT) versions 21.11.6 and 22.10.7 for CDC317 and Cp14 respectively. Both runs were set to the default fast configurations. Basecalling and demultiplexing was run within MinKNOW during sequencing. This generated 638,855 and 237,020 raw reads respectively for CDC317 and Cp14.

### Sequence analysis

The Illumina reads for all 42 *C. parapsilosis* samples were downsampled to ∼100x coverage using the Picard version 2.21.2 DownsampleSam on unmapped SAM files. These files were converted to FASTQ format using Picard SamToFastq and aligned to the *C. parapsilosis* CDC317 reference genome using bwa mem version 0.7.17 (65). GATK version 4.1.4.1 was used to mark duplicate reads and reorder the mapped BAM files with the tools MarkDuplicates and ReorderSam respectively (66). Variants were called on individual samples using GATK HaplotypeCaller with the “-ERC GVCF” tag. The GVCF outputs were combined into a multi-sample VCF with GATK CombineGVCFs, and then genotyped using GATK GenotypeGVCFs. SNP variants were hard filtered using GATK VariantFiltration with following parameters: QD < 2.0, QUAL < 30.0, SOR > 3.0, FS > 60.0, MQ < 40.0, MQRankSum < -12.5, ReadPosRankSum < -8.0 and DP < 10. Indels were similarly filtered using parameters: QD < 2.0, QUAL < 30.0, FS > 200.0, and ReadPosRankSum < -20.0. All variants were filtered using GATK SelectVariants to remove multi-allelic sites, and sites that contained >10% ‘no-call’ genotypes. The two sets of variants were combined for GWAS analysis, and a file containing only SNPs was used for phylogenetic analysis. MinION reads for isolate CDC317 were filtered using NanoFilt to exclude reads with quality < 12 and length < 10000 (67). The filtered reads were assembled using Canu version 2.2 (68). Errors in this assembly were removed by incorporating Illumina read data using NextPolish version 1.4.0 (69). The assembly is available under accession PRJNA1031570. Reads for *C. orthopsilosis* were obtained from Schröder et al. (70), Pryszcz et al. (71), Bergin et al. (44) and Zhai et al. (63), and reads for *C. metapsilosis* from Zhai et al. (63) and O’Brien et al. (72) (Table S2).

### Phylogeny

A FASTA alignment of all sites containing a SNP in at least one isolate was created from the multi-sample VCF file using a custom script (https://github.com/CMOTsean/HetSiteRando). Heterozygous variants were randomly assigned to either allele on a per-site basis. A SNP tree was constructed with the alignment file using RAxML version v8.2.12 with the GTRGAMMA model of nucleotide substitution and 1000 bootstrap replicants (73).

### GWAS

A binary phenotype matrix where all samples were scored as either resistant or susceptible was used as input to the GWAS. Variants entering the GWAS were filtered to only those likely to affect protein function, as annotated by SIFT4G (74). PLINK version 1.90b6.21 was used to reformat input data into BED, BIM, and FAM files for the GWAS (75). GEMMA version 0.98.5 was used to create a relatedness matrix between all strains, and then to conduct the GWAS itself using parameters ‘-hwe 0.0 -maf 0.0’ (51).

### Estimating repeat copy number and structure

To calculate the estimated copy number of the *CPAR2_304370* locus, the average coverage across the ORF in each isolate was found using BEDTools coverage version 2.29.2 and then divided by half the modal genome coverage (BEDTools genomecov) for that isolate (76).

To confirm copy number estimates and investigate the structure of the repeat, the nucleotide sequence of *CDR1B.1* was searched against MinION reads from strains CDC317 and Cp14 using BLASTN. Additionally, the Cp14 reads were aligned to the MinION CDC317 assembly using GraphMap version 0.3.0 (77).

### *C. parapsilosis* species complex comparison

To create the synteny maps between *C. parapsilosis* and its sister species, the *C. parapsilosis* CDC317 reference genome was aligned against the *C. metapsilosis* BP57 reference genome (GCA_017655625.1) (52) and the *C. orthopsilosis* SY36 long-read assembly (PRJNA767198) using BLASTN. Hits from each query chromosome were assigned a colour and then plotted.

To construct the tree of CDR protein sequences, the sequences for *C. albicans*, *C. parapsilosis*, and *C. metapsilosis* were taken from CGOB with the exception of *CMET_1535* which came from the *C. metapsilosis* BP57 assembly.

## Data availability

DNA sequence assembly and raw data is available under accession PRJNA1031570 and RNA sequencing is available at BioProject 14022043.

## Acknowledgements

This work was supported by Science Foundation Ireland (grant numbers 19/FFP/6668 and 18/CRT/6214 to G.B. and 20/FFP-A/8795 to K.H.W.) and by NIAID grant U19AI110818 to the Broad Institute (C.A.C.). This work was also supported by NIH NIAID grant R01 A1058145 awarded to P.D.R. and in part by the National Cancer Institute of the National Institutes of Health under Award Number P30 CA021765 awarded to the Hartwell Center at St. Jude Children’s Research Hospital. We thank Daniel J. Diekema at the University of Iowa for generously providing the clinical isolates used in this study. We are grateful to Adam Ryan, Eoin Ó Cinnéide, Athaliah Fubara and Conor Hession in UCD for help with MinION sequencing of Cp14 and CDC317, and Evelyn Zuniga for some MIC assays.

## Supplementary Material

**Table S1:** GEMMA (GWAS) analysis

**Table S2.** Sequencing data for isolates.

| Isolate name | City | Country | Sequence data source | Reference |
| --- | --- | --- | --- | --- |
| **Candida parapsilosis** |  |  |  |  |
| Cp1 | Bloemfontein | South Africa | This study | (40) |
| Cp10 | Bloemfontein | South Africa | This study | (40) |
| Cp11 | Bloemfontein | South Africa | This study | (40) |
| Cp12 | Bloemfontein | South Africa | This study | (40) |
| Cp13 | New York | United States | This study | (40) |
| Cp14 | Helsinki | Finland | This study | (40) |
| Cp15 | Quito | Ecuador | This study | (40) |
| Cp16 | Bloemfontein | South Africa | This study | (40) |
| Cp17 | Bloemfontein | South Africa | This study | (40) |
| Cp18 | Bloemfontein | South Africa | This study | (40) |
| Cp19 | Bloemfontein | South Africa | This study | (40) |
| Cp2 | Bloemfontein | South Africa | This study | (40) |
| Cp20 | Bloemfontein | South Africa | This study | (40) |
| Cp21 | Ann Arbor | United States | This study | (40) |
| Cp22 | Johannesburg | South Africa | This study | (40) |
| Cp24 | Johannesburg | South Africa | This study | (40) |
| Cp25 | Johannesburg | South Africa | This study | (40) |
| Cp26 | Johannesburg | South Africa | This study | (40) |
| Cp27 | Hershey | United States | This study | (40) |
| Cp28 | Bloemfontein | South Africa | This study | (40) |
| Cp29 | Bloemfontein | South Africa | This study | (40) |
| Cp3 | Kuala Lumpur | Malaysia | This study | (40) |
| Cp30 | Bratislava | Slovakia | This study | (40) |
| Cp31 | Johannesburg | South Africa | This study | (40) |
| Cp32 | Caracas | Venezuela | This study | (40) |
| Cp34 | Johannesburg | South Africa | This study | (40) |
| Cp35 | Johannesburg | South Africa | This study | (40) |
| Cp36 | Detroit | United States | This study | (40) |
| Cp37 | Johannesburg | South Africa | This study | (40) |
| Cp38 | Johannesburg | South Africa | This study | (40) |
| Cp39 | Bratislava | Slovakia | This study | (40) |
| Cp4 | Johannesburg | South Africa | This study | (40) |
| Cp5 | New York | United States | This study | (40) |
| Cp6 | Turino | Italy | This study | (40) |
| Cp7 | Johannesburg | South Africa | This study | (40) |
| Cp8 | Bloemfontein | South Africa | This study | (40) |
| Cp9 | Bloemfontein | South Africa | This study | (40) |
| CDC317 | Mississippi | USA | PRJNA795920 | (79) |
| FM16 | Nantes | France | PRJNA795920 | (44) |
| MSK809 | New York | USA | PRJNA748054 | (44) |
| 73-037 | Leeds | UK | PRJNA795920 | (44) |
| 73-114 | Leeds | UK | PRJNA795920 | (44) |
| **Candida orthopsilosis** |  |  |  |  |
| 151 | Atlanta | USA | PRJNA322245 | (70) |
| 185 | Atlanta | USA | PRJNA322245 | (70) |
| 282 | Baltimore | USA | PRJNA322245 | (70) |
| 320 | Baltimore | USA | PRJNA322245 | (70) |
| 421 | Pisa | Italy | PRJNA322245 | (70) |
| 422 | Pisa | Italy | PRJNA322245 | (70) |
| 423 | L'Aquila | Italy | PRJNA322245 | (70) |
| 424 | Pisa | Italy | PRJNA322245 | (70) |
| 425 | Pisa | Italy | PRJNA322245 | (70) |
| 426 | Varese | Italy | PRJNA322245 | (70) |
| 427 | Pisa | Italy | PRJNA322245 | (70) |
| 428 | Hong Kong | Hong Kong | PRJNA322245 | (70) |
| 433 | Sint-Niklaas | Belgium | PRJNA322245 | (70) |
| 434 | Bristol | UK | PRJNA322245 | (70) |
| 435 | Pisa | Italy | PRJNA322245 | (70) |
| 436 | Pisa | Italy | PRJNA322245 | (70) |
| 437 | Pisa | Italy | PRJNA322245 | (70) |
| 498 | Baltimore | USA | PRJNA322245 | (70) |
| 504 | Baltimore | USA | PRJNA322245 | (70) |
| 599 | Baltimore | USA | PRJNA322245 | (70) |
| 748 | Atlanta | USA | PRJNA322245 | (70) |
| 831 | Atlanta | USA | PRJNA322245 | (70) |
| 1540 | Baltimore | USA | PRJNA322245 | (70) |
| 1799 | Atlanta | USA | PRJNA322245 | (70) |
| 1825 | Baltimore | USA | PRJNA322245 | (70) |
| 90-125 | N/A | N/A | PRJNA431439 | (70) |
| B-8274 | N/A | Pakistan | PRJNA322245 | (70) |
| B-8323 | N/A | Pakistan | PRJNA322245 | (70) |
| MCO456 | N/A | N/A | PRJEB4430 | (71) |
| MSK477 | New York | USA | PRJNA579121 | (63) |
| MSK479 | New York | USA | PRJNA579121 | (63) |
| MSK616 | New York | USA | PRJNA748054 | (44) |
| MSK636 | New York | USA | PRJNA579121 | (63) |
| MSK638 | New York | USA | PRJNA579121 | (63) |
| MSK639 | New York | USA | PRJNA579121 | (63) |
| MSK805 | New York | USA | PRJNA748054 | (44) |
| **Candida metapsilosis** |  |  |  |  |
| MSK403 | New York | USA | PRJNA579121 | (63) |
| MSK404 | New York | USA | PRJNA579121 | (63) |
| MSK413 | New York | USA | PRJNA579121 | (63) |
| MSK414 | New York | USA | PRJNA748054 | (72) |
| MSK415 | New York | USA | PRJNA579121 | (63) |
| MSK416 | New York | USA | PRJNA579121 | (63) |
| MSK417 | New York | USA | PRJNA579121 | (63) |
| MSK418 | New York | USA | PRJNA579121 | (63) |
| MSK429 | New York | USA | PRJNA579121 | (63) |
| MSK430 | New York | USA | PRJNA579121 | (63) |
| MSK431 | New York | USA | PRJNA579121 | (63) |
| MSK432 | New York | USA | PRJNA579121 | (63) |
| MSK433 | New York | USA | PRJNA579121 | (63) |
| MSK434 | New York | USA | PRJNA579121 | (63) |
| MSK445 | New York | USA | PRJNA579121 | (63) |
| MSK446 | New York | USA | PRJNA579121 | (63) |
| MSK447 | New York | USA | PRJNA579121 | (63) |
| MSK448 | New York | USA | PRJNA579121 | (63) |
| MSK449 | New York | USA | PRJNA579121 | (63) |
| MSK450 | New York | USA | PRJNA579121 | (63) |
| MSK461 | New York | USA | PRJNA579121 | (63) |
| MSK462 | New York | USA | PRJNA579121 | (63) |
| MSK463 | New York | USA | PRJNA579121 | (63) |
| MSK464 | New York | USA | PRJNA579121 | (63) |
| MSK465 | New York | USA | PRJNA579121 | (63) |
| MSK466 | New York | USA | PRJNA579121 | (63) |
| MSK606 | New York | USA | PRJNA748054 | (72) |
| MSK607 | New York | USA | PRJNA748054 | (72) |
| MSK798 | New York | USA | PRJNA748054 | (72) |
| MSK801 | New York | USA | PRJNA748054 | (72) |

**Figure S1.**
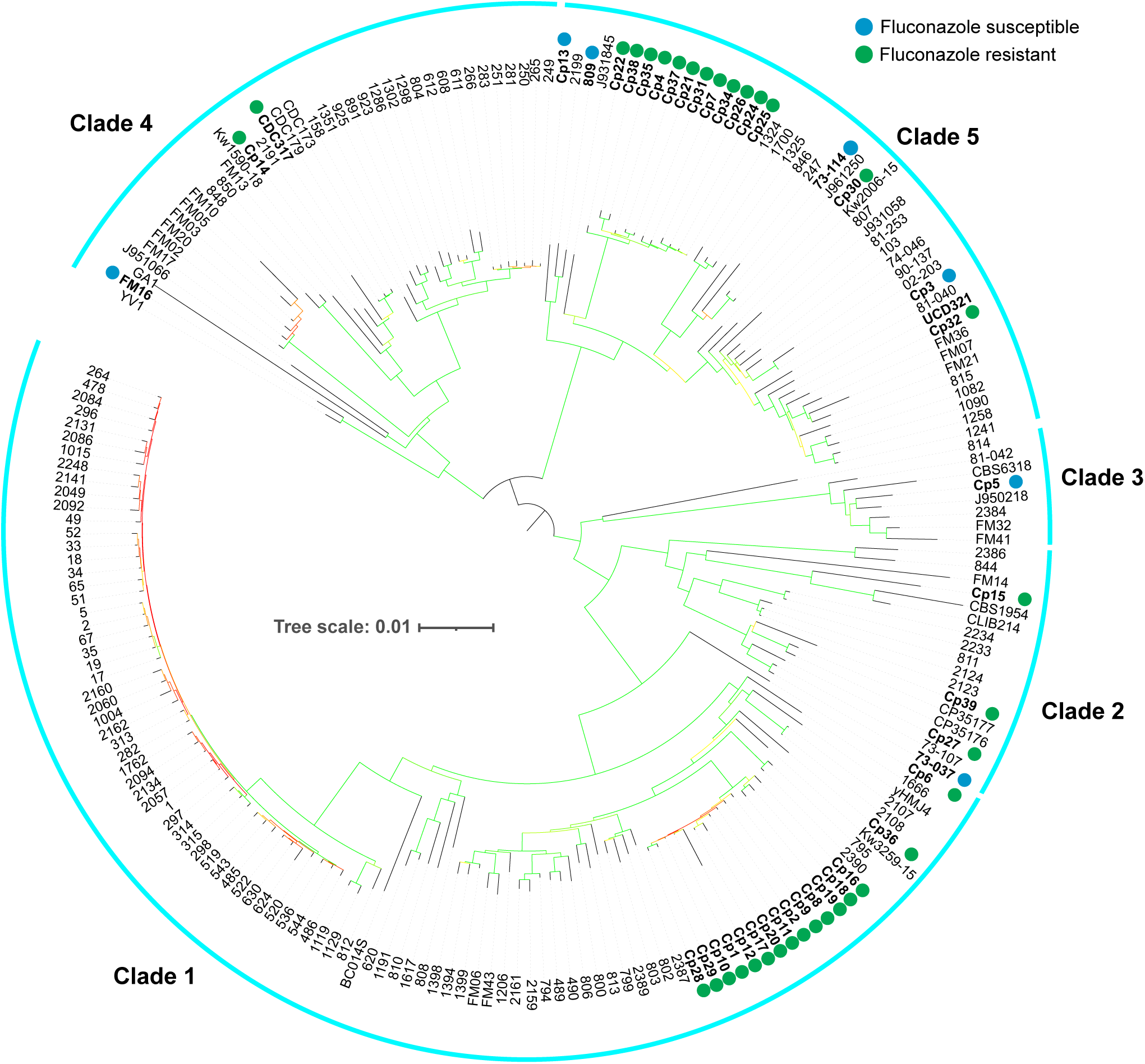
Maximum-likelihood tree of 207 *Candida parapsilosis* isolates constructed as in Figure 1. Additional isolates, and clade designations were taken from (44). Susceptible isolates in this paper are marked with a blue circle, and resistant isolates are marked with a green circle. Coloured branches indicate bootstrap values after 1000 iterations of bootstrap sampling for that branch ranging from 0 (red) to 100 (green).

**Figure S2.**
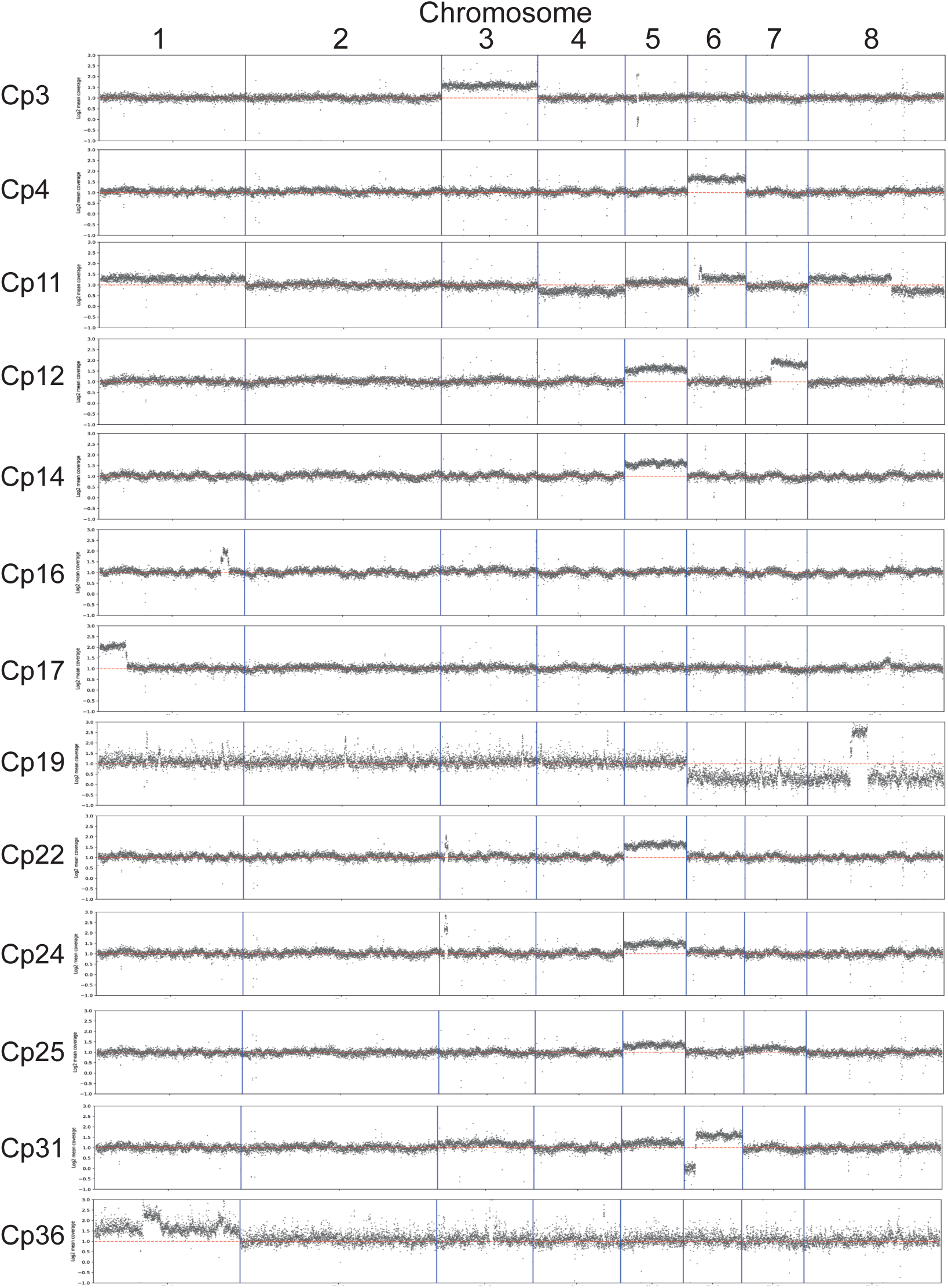
Genome-wide coverage tracks for aneuploid strains. Coverage was calculated as the log2 mean coverage of 1kb windows and plotted. Dashed red line shows log2 value = 1, i.e. typical 2x sdiploid coverage

